# Correcting differential gene expression analysis for cyto-architectural alterations in substantia nigra of Parkinson’s disease patients reveals known and potential novel disease-associated genes and pathways

**DOI:** 10.1101/2021.05.25.445590

**Authors:** Ferraro Federico, Fevga Christina, Bonifati Vincenzo, Mandemakers Wim, Mahfouz Ahmed, Reinders Marcel

**Affiliations:** Department of Clinical Genetics, Erasmus MC, University Medical Center Rotterdam, Rotterdam, the Netherlands; Delft Bioinformatics Lab, Delft University of Technology, Delft, the Netherlands; Leiden Computational Biology Center, Leiden, the Netherlands; Department of Biomedical Data Sciences, Section Molecular Epidemiology, Leiden University Medical Center, Leiden, the Netherlands; Department of Human Genetics, Leiden University Medical Center, Leiden, the Netherlands

## Abstract

Several studies have analyzed gene expression profiles in the substantia nigra to better understand the pathological mechanisms causing Parkinson’s disease (PD). However, the concordance between the identified gene signatures in these individual studies was generally low. This might be caused by a change in cell type composition as loss of dopaminergic neurons in the substantia nigra pars compacta is a hallmark of PD. Through an extensive meta-analysis of nine previously published microarray studies, we demonstrated that a big proportion of the detected differentially expressed genes was indeed caused by cyto-architectural alterations due to the heterogeneity in the neurodegenerative stage and/or technical artifacts. After correcting for cell composition, we identified a common signature that deregulated the previously unreported ammonium transport, as well as known biological processes including bioenergetic pathways, response to proteotoxic stress, and immune response. By integrating with protein-interaction data, we shortlisted a set of key genes, such as *LRRK2, PINK1*, and *PRKN* known to be related to PD; others with compelling evidence for their role in neurodegeneration, such as GSK3β, WWOX, and VPC; as well as novel potential players in the PD pathogenesis, including *NTRK1, TRIM25, ELAVL1*. Together, these data showed the importance of accounting for cyto-architecture in these analyses and highlight the contribution of multiple cell types and novel processes to PD pathology providing potential new targets for drug development.

**Significance Statement:** The exploration of the transcriptomic landscape in PD is pivotal for the understanding of the pathological mechanisms of this disease. Nonetheless, little attention has been paid to the influence of cell composition on the transcriptome even though it is known that cyto-architecture undergoes major alterations in neurodegenerative diseases such as PD. Our study signifies that changes in cellular architecture of human *substantia nigra* in PD have a strong effect on the set of detected differentially expressed genes. By reanalyzing the data and accounting for cell composition, we provide an updated description of deregulated biological processes in PD and nominate a shortlist of PD-associated genes for further investigations.

## Introduction

Parkinson’s disease (PD) is the second most common neurodegenerative disorder after Alzheimer’s disease. In PD, the loss of dopaminergic neurons in the *substantia nigra* pars compacta and neurodegeneration in other brain areas leads to motor as well as non-motor manifestations (Surmeier et al., 2017; Poewe et al., 2017; Balestrino and Schapira, 2020). Alpha-synuclein positive inclusions, termed Lewy bodies (LB) and Lewy neurites, are found in the surviving neurons (Fares et al., 2021). Despite the elusiviness of the biogenesis and spreading of these structures, according to Braak’s model (Braak et al., 2003), LB-pathology spreads in the PD brain along a caudo-rostral trajectory, correlating with disease progression.

Notwithstanding great progress since its initial description (Parkinson, 1817), the causative factors remain poorly understood. Various environmental, lifestyle, and genetic factors, including rare and highly penetrant variants in a limited number of genes (Bandres-Ciga et al., 2020) and 90 common risk loci (Nalls et al., 2019), have been implicated in its pathogenesis. Still, a big proportion of missing heritability remains.

In parallel to genetic studies, genome-wide expression profiling has been used to characterize alterations in molecular pathways in different brain regions, blood, and other tissues of PD patients (Borrageiro et al., 2018). For example, a recent transcriptomics study reported evidence of differential brain regional vulnerability to PD progression in accordance with the Braak’s hypothesis (Keo et al., 2020). Based on its role in PD, the *substantia nigra*, has been extensively investigated in these genome-wide expression profiling studies.

Further insight into PD pathogenic pathways has been enabled by the aggregation of small-scale, low-powered, and low concordance studies (Oerton et al., 2017) into larger meta-analyses, which has led to the nomination of putative key regulators of disease progression (Zheng et al., 2010; Wang et al., 2019a). This approach has proven to be fruitful, even in the context of a high degree of heterogeneity in the putative causes and severity of phenotypes of the included patients.

Unfortunately, transcriptomic studies suffer from various technical limitations, such as RNA degradation, that affect the different cell types to a variable extent (Jaffe et al., 2017). Furthermore, differences in cyto-architecture originating from biological heterogeneity (due to e.g. age, gender, genetic background), as well as sample preparation can further influence downstream analyses. An even stronger confounder might be represented by cell composition changes induced by a pathology (Srinivasan et al., 2016). This issue is particularly concerning since it is not possible to distinguish between changes in genes that are highly expressed in a cell type whose proportion change and a genuine pathology-related transcriptional deregulation. While the former might be interesting as it may correlate with disease progression, the latter can inform on the molecular mechanism for which therapeutic strategies might be devised. In neurodegenerative disorders and especially in highly affected brain regions, as the *substantia nigra* in PD, this phenomenon might be even more pronounced.

To tackle these and other challenges, single cell RNA sequencing (scRNAseq) and single nucleus RNA sequencing (snRNAseq) are currently rising in popularity. However, thus far only a limited number of PD substantia nigra samples have been profiled at the single cell level (Smajić et al., 2021) severely decreasing the power to detect relevant differences. Alternatively, cell proportions can be estimated from bulk transcriptomics data, and then analyses can be corrected for altered cyto-architecture. Recently, several bioinformatic approaches have been proposed to estimate and use these surrogates for the proportions of the cell types, offering the opportunity to exploit the enormous amount of data readily available in public repositories (Shen-Orr et al., 2010; Mancarci et al., 2017; McKenzie et al., 2018; Newman et al., 2019; Wang et al., 2019b; Zaitsev et al., 2019; Dong et al., 2020; Li et al., 2020).

In this study, we systematically assessed the transcriptomic evidence for the presence of changes in cell compositions in expression data of nine publicly available Parkinson’s related microarrays of the *substantia nigra* (see Figure 1 for an overview). We conducted the first meta-analysis of these data sets while evaluating the impact of the cyto-architectural alterations on the differential expression analysis. By correcting for these effects, we were able to detect genuine disease-related changes in the transcriptional landscape of the *substantia nigra* in PD patients. Finally, we explored the protein interactome of the identified deregulated genes and nominated promising candidates for further investigations by exploiting their network characteristics.

**Fig.1.**
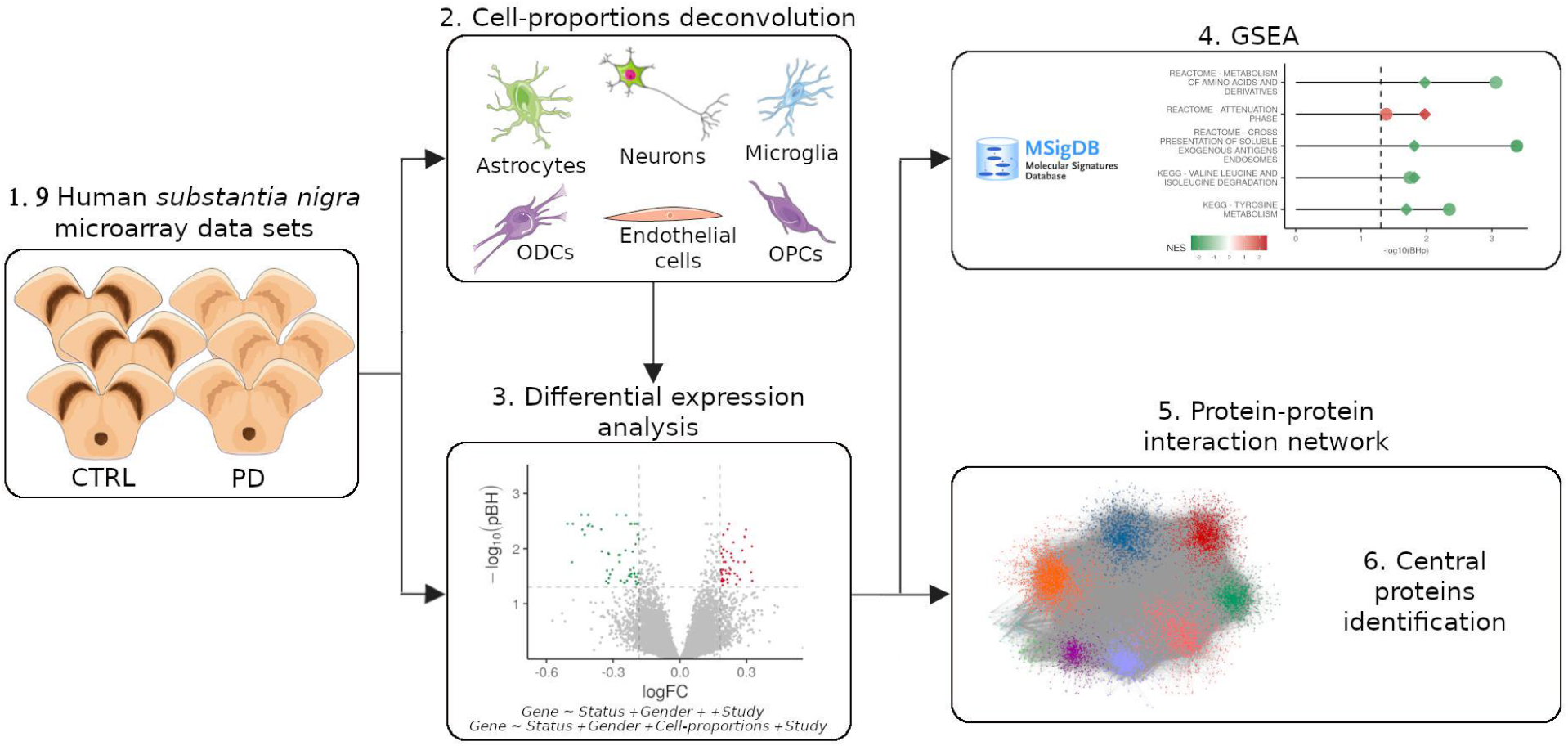
Summary of this study. **1**. Human substantia nigra microarrays from PD and CTRL were downloaded from GEO. **2**. Cell-proportion deconvolution for six cell types. **3**. Differential expression analysis with and without cell proportions. **4**. GSEA on the expression matrix. **5**. Central protein identification in PPI network around detected differentially expressed genes. Drawing of substantia nigra and cells were obtained from Servier Medical Art templates (Creative Commons Attribution 3.0 Unported License; https://smart.servier.com).

## Materials and Methods

### 1. Transcriptome data sets download, pre-processing, and gender imputation

We downloaded gene expression data sets from the Gene Expression Omnibus (GEO) using “Parkinson’s disease” and “*substantia nigra*” as keywords. In total we found 10 studies: nine (GSE7621, GSE20333, GSE20292, GSE20163, GSE20164, GSE54282, GSE49036, GSE43490, and GSE42966) analyzed the whole *substantia nigra* for each patient; one analyzed separately the medial and the lateral part of the *substantia nigra* (GSE8397) from which we only used the lateral as more affected in PD (Duke et al., 2007). GSE54282 was excluded from analysis because of the small number of available samples. Only probes mapping to an Entrez ID using the biomaRt package version 2.42.1 (Durinck et al., 2005) were kept. When multiple probes mapped to the same Entrez ID, we kept the one with the maximum variance and connectivity using the *collapseRows* function from the WGCNA package version 1.69 (Miller et al., 2011), resulting in a total of 18,948 genes. We used gene expression as already processed in the original studies (either Robust Multichip Average normalization or the Affymetrix microarray suite), removed outlier samples according to the original publications. Finally, expressions were log transformed, the merged data set was quantile normalized with the *normalizeBetweenArrays* function from the limma package version 3.42.2 (Ritchie et al., 2015). Since gender information was not always available, we used the massiR package version 1.22.0 (Buckberry et al., 2014) to annotate all the samples using the top 75% variable genes for the prediction.

### 2. DEGs meta-analysis

The DEGs were identified by fitting a linear mixed-effects model (LMM). For each gene, two LMMs were fitted using the *lmer* function from the lmerTest package version 3.1.2. One LMM accounts for the status and the gender as fixed effects and the different studies as random-effects: *Gene*_*exp*_ *∼ Status + Gender + (1*|*Study)*, with *(1*|*Study)* indicating a one-hot encoding of the study. The other LMM, accounts also for estimates of cell types (Neurons (NEU), Oligodendrocytes (ODC) and oligodendrocytes precursor cells (OPC)) as fixed effects: *Gene*_*exp*_ *∼ Status + NEU + ODC + OPC + Gender+(1*|*Study)*. We chose to only use a subset of the cell types as correction covariates to avoid the introduction of collinear predictors. Specifically, we included those cell types whose estimates are only weakly correlated with the estimates of the other cell types in the model but highly correlated with the excluded ones (Extended Data Figure 1-1B). Moreover, we ensured to include cell types whose estimates show changes in opposite directions between PD and CTRL (Figure 1B). P-values were corrected using the Benjamini-Hochberg (BH) method.

### 3. Cell type markers identification from *Substantia nigra* single cell data

Human *substantia nigra* single cell data (Agarwal et al., 2020) was downloaded from GEO (GSE140231) and preprocessed with Seurat package version 2.3.0 (Butler et al., 2018) according to the original publication. Cell markers for each cluster compared to all the others were identified from transcripts detected in at least 30% of the available cells, and a log Fold Change higher than 0.5 using the function *FindAllMarkers*. The clusters were broadly annotated into six cell types using well-known markers: *GFAP* and *GINS3* for the astrocytes; *MOBP* and *MOG* for the oligodendrocytes; *CSF1R* and *OLR1* for the microglia; *GAD1, GAD2, GABRA*, and *TH* for the neurons; *VCAN* for the oligodendrocyte precursor cells; and *LGALS1* and *RGS5* for the endothelial cells.

### 4. Cell-type proportions estimation and marker selection

Cell type proportions were estimated by deconvolution using the function *MDeconv* from the TOAST package version 1.1.2 (Li et al., 2019; Li et al., 2020). We considered six brain cell types: astrocytes, endothelial cells, neurons, microglia, ODCs and OPCs. As cell markers, we initially tested 1) the set of markers identified from the s*ubstantia nigra* single cell data (GSE140231), and 2) the set of 5,500 markers obtained from the Brain Cell Type Specific Gene Expression Analysis package version 1.0.0 (BRETIGEA, McKenzie et al., 2018). To define which of the two sets, the number, and identity of the markers to employ, we adopted a quality-based heuristic selection. For each marker set, each cell type and study, we ranked the markers on the basis of their association to the 1^st^ principal component. Next, we calculated the Pearson’s correlation among the top 100, and iteratively excluded those showing negative correlations with the majority of the others until no markers with opposite correlations were left. Finally, we selected the top 20 markers for each cell type. We decided to use the BRETIGEA-derived markers in the rest of the analyses since they resolved better into clusters in the correlation matrices (Extended Data Figure 1-2).

### 5. Cell-types proportions comparison and statistical analyses

For each study and cell type, statistically significant differences in the estimates of a cell type proportion between the CTRL and the PD *substantia nigra* were assessed with a linear model controlling for the other annotated covariates, i.e. *Proportion*_*CellType*_ *∼ Status + Gender + Age + BraakStaging*. We also conducted a meta-analysis to verify the alterations in proportion across the studies. To this aim, a random-effects meta-analysis was conducted in metafor version 2.1.0 (Viechtbauer et al., 2010). For each cell type, the standardized mean difference was calculated, and the associated p-values were corrected by the BH method.

### 6. Gene set enrichment analysis

Gene set enrichment analysis (GSEA) was performed using the Fast gene set enrichment analysis package version 1.12.0 (fgsea, Korotkevich et al., 2019), 100,000 permutations, with the Gene Ontology (GO) and the Canonical data set downloaded from the Molecular Signatures Database (MSigDB). For the GSEA on the gene expression, the genes in our data set were first ranked in descending order by the negative logarithm in base 10 of the adjusted p-values multiplied for the sign of the effect size. For the GSEA of the PPI network, nodes were ranked by the betweenness centrality (Zito et al., 2021). All p-values were corrected using the BH method.

### 7. Expression-weighted cell-type enrichment (EWCE)

Gene set enrichment for specific cell types was done using the expression-weighted cell-type enrichment (EWCE, Skene and Grant, 2016) package version 0.99.2. To this aim, we first calculated the average expression matrix for each cell type in the *substantia nigra* using the *AverageExpression* from the Seurat package using the GSE140231 data set. For each tested list, 10.000 randomly sampled (equal sized) gene sets from all genes in the average expression matrix were used to calculate the p-values, then adjusted by the BH correction.

### 8. Protein-protein-interaction network construction and analysis

A protein-protein interaction network was built from six publicly available databases: 1) the Human Reference Interactome and Literature Benchmark (HuRI, Luck et al., 2020), 2) the Biological General Repository for Interaction Datasets (BioGRID build 3.5.186, Oughtred et al., 2019), 3) STRING v11 database (Szklarczyk et al., 2019), 4) the Integrated Interactions Database, Version 2018-11 (IID, Kotlyar et al., 2018), 5) BioPlex 3.0 (Huttlin et al., 2020), and 6) IntAct Database (Orchard et al., 2014). For all databases, only experimentally validated interactions were selected, the lists were merged, non-human proteins, nodes with just one edge, self-interactions, and duplicated interactions were removed. The final graph was visualized in Cytoscape 3.8.0 and analyzed with the R package igraph 1.2.5 (Csardi et al., 2006). Fisher’s exact test (FET) was used to test for the enrichment in the whole network and in the central proteins for typical PD-causing genes, selected according to MDSGene (https://www.mdsgene.org/g4d), and for the genes in proximity of PD-risk variants (Nalls et al., 2019, downloaded from GWAS Catalog, https://www.ebi.ac.uk/gwas/). The number of human protein-coding genes was used as background for the tests on the whole network, while the network size was used for the ones with the central proteins.

### 9. Code and data accessibility

All analyses were conducted in R version 3.6.3 on a laptop (Intel^R^ Core™ i7-7700HQ CPU 2.80GHz, 8 Gb RAM) running Linux Ubuntu 18.04. All code is freely available online at https://github.com/f-ferraro/CytoarchitecturePDsn. Data sets analyzed can be freely downloaded from https://www.ncbi.nlm.nih.gov/geo/ using the accession codes GSE7621, GSE8397, GSE20333, GSE20292, GSE20163, GSE20164, GSE49036, GSE43490, GSE42966, and GSE140231.

## Results

### 1. Alterations in composition of the *substantia nigra* are heterogeneous

In total, 70 control (CTRL) and 88 PD transcriptomes were put together in our study. After imputing the gender for all the samples (Methods), there were 62 females (27 CTRL, 35 PD) and 96 males (43 CTRL, 53 PD), with an average age of death of 75 years (Table 1). To capture the PD induced cyto-architectural alterations, we estimated the proportions of six cell types (astrocytes, endothelial cells, neurons, microglia, ODCs, and OPCs) from the bulk data using computational deconvolution (Methods). This revealed cell type heterogeneity; not only between the two groups, i.e. CTRL and PD patients, but also within each group (across samples), as well as between data sets (Figure 2A, Extended Data Table 2-1). Although we observed similar trends between CTRL and PD brains, the alterations in cell proportions did not consistently replicate across the analyzed data sets. To identify consistent alterations, shared among data sets, we conducted a meta-analysis of the estimated cell type proportions. This showed a significant increase in endothelial cells and oligodendrocytes as opposed to a decrease in the neuronal estimates (Figure 2B, Extended Data Table 2-2). The change in neurons was the strongest, followed by the ODCs and then endothelial cells.

**Table 1:** Overview of the studies contributing data to our analyses.

| GEO ID | Tissue | #CTRL<br>(F:M) | #PD<br>(F:M) | Platform | #N<br>Probes | F:M | Average<br>age of<br>death | Reference |
| --- | --- | --- | --- | --- | --- | --- | --- | --- |
| GSE7621 | SN | 9 (5:4) | 16 (3:13) | GPL570 | 54675 | 8:17 | 78 | Lesnick et al., 2007 |
| GSE8397 | LatSN | 7 (2:5) | 9 (4:5) | GPL96 | 22283 | 6:10 | 76 | Moran et al., 2006 |
| GSE20333 | SN | 6 (1:5) | 6 (5:1) | GPL201 | 8793 | 6:6 | 78 | Edna et al., 2010 |
| GSE20292 | SN | 15 (5:10) | 11 (6:5) | GPL96 | 22283 | 11:15 | 73 | Zheng et al., 2010 |
| GSE20163 | SN | 9 (1:8) | 8 (4:4) | GPL96 | 22283 | 5:12 | 74 | Zheng et al., 2010 |
| GSE20164 | SN | 5 (4:1) | 6 (2:4) | GPL96 | 22283 | 6:5 | 82 | Zheng et al., 2010 |
| GSE49036 | SN | 8 (5:3) | 15 (7:8) | GPL570 | 54675 | 12:11 | 78 | Dijkstra et al., 2015 |
| GSE43490 | SN | 5 (2:3) | 8 (0:8) | GPL6480 | 41108 | 2:11 | 74 | Corradini et al., 2014 |
| GSE42966 | SN | 6 (2:4) | 9 (4:5) | GPL4133 | 45220 | 6:9 | 73 | Bando et al., 2018 |

**Figure 2:**
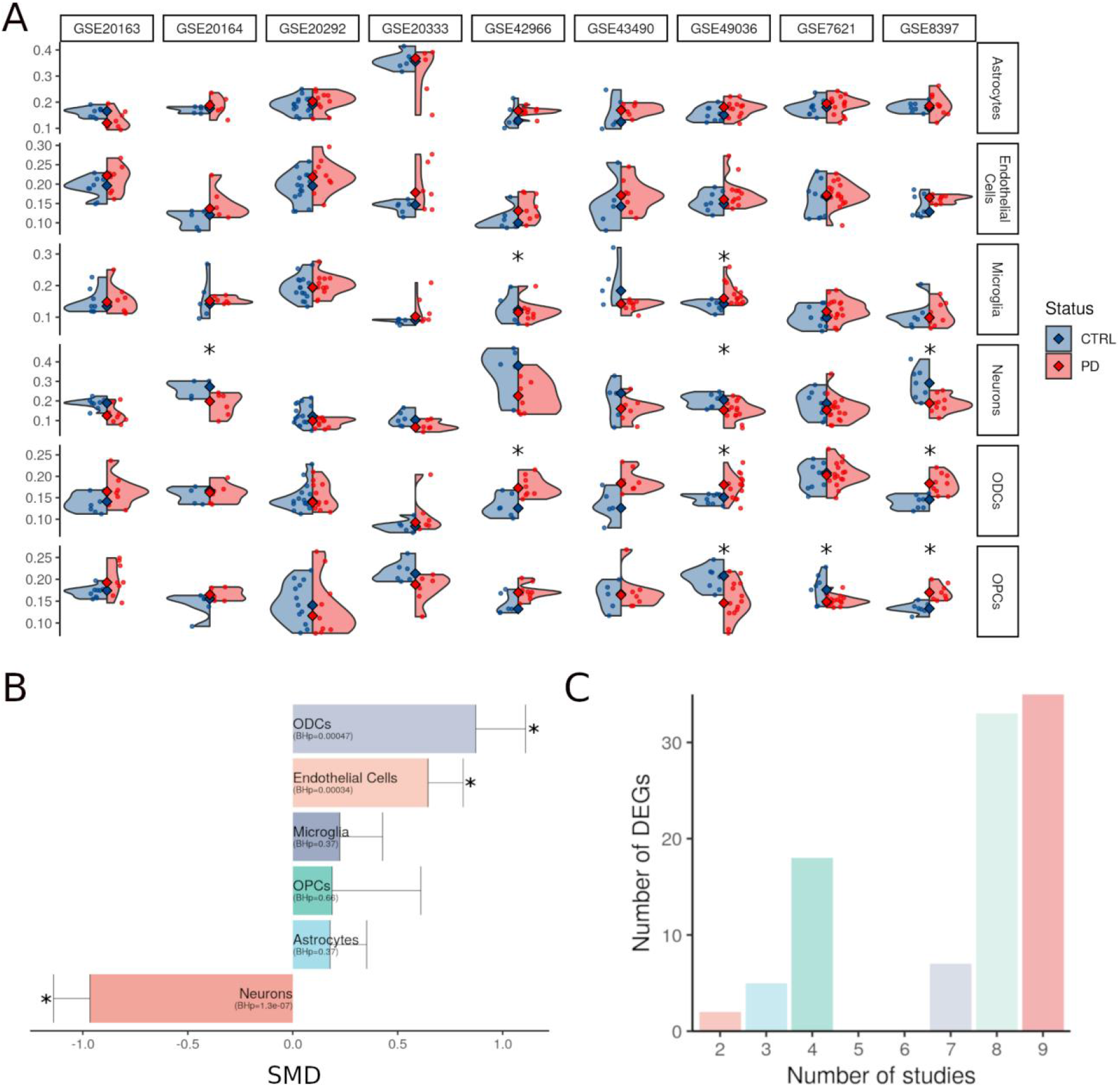
Cyto-architectural heterogeneity as estimated from bulk transcriptomics using the deconvolution strategy of TOAST. **A**. Cell estimates of six cell-types in substantia nigra across different data sets and conditions. Blue CTRL, red PD. Significant variations are indicated by asterisk (p<0.05) **B**. Bar plot of the standardized mean differences (SMD) in cell-estimates from the random-effects meta-analysis conducted with metafor. BH-adjusted p-values are reported between brackets, significant differences are indicated by asterisk (BHp<0.05), and standard error for each cell type is reported as error bar. **C**. Number of DEGs of the cell-proportions-aware LMM probed by a specific number of microarrays in our data set.

### 2. The cyto-architectural alterations are a major confounder in the DEGs identification

To detect consistent differentially expressed genes (DEGs), we fitted a linear mixed-effects model (LMM) for each gene across all data sets, with a random-effect for the different studies and fixed effects for gender and CTRL/PD status (Methods). Out of the 1,072 deregulated genes between PD and control samples (adjusted p-value <0.05, fold change (FC)> 1.2), 713 were down-regulated and 359 were up-regulated (Extended Data Figure 1-1A, Extended Data Table 1-3). We then included estimates of the neurons, ODCs, and OPCs as fixed effects into the linear mixed effects model (Methods). We chose to correct only for these three cell types due to their collinearity with the other cell types (Extended Data Figure 1-1B). With this second LMM, we detected a reduced number of DEGs (adjusted p-value <0.05, FC> 1.2): 100 instead of the 1,072 obtained with the cell-proportions-unaware LMM (67 genes overlapping). Out of the 100, 55 were down-regulated and 45 were up-regulated in PD with respect to the CTRL samples (Extended Data Table 2-3, Extended Data Figure 2-1B). The majority of the DEGs (68%) was probed in more than seven studies, showing a robust alteration signature across the data sets despite possible heterogeneity in etiological factors and pathology severity (Figure 2D). None of the PD-causing genes nor any of the genes proximal to GWAS risk variants were found among the DEGs. To find out for which cell types the 100 DEGs are enriched, we performed an expression-weighted cell-type enrichment (EWCE) (Methods). Down-regulated genes appeared to be significantly enriched in neurons, whereas up-regulated genes were instead enriched in OPCs (Extended Data Table 2-4, Extended Data Figure 2-1C). No significant enrichment was found for the other cell types.

### 3. GSEA reveals that the majority of the altered pathways in PD are down-regulated

To explore the effect of the PD-related gene expression deregulation, we conducted a gene set enrichment analysis (GSEA) for functions in GO and the MSigDB canonical data set (Methods). Genes were ranked by the signed corrected -log10 p-values obtained from either LMM. Similarly, to the DEG analyses, accounting for cell composition decreased the number of significant pathways (adjusted p-value <0.05) (Figure 3, Extended Data Table 3-1). Specifically, 258 significant canonical pathways (94 up-regulated and 164 down-regulated) were identified with the cell-proportions-unaware (first) LMM. When applied on the expression matrix ranked by the cell-proportion aware (second) LMM, only 9 pathways (1 up-regulated and 8 down-regulated) remained significant, and 1 pathway, the up-regulation of the Heat Shock Transcription Factor 1 (*HSF1*) activation pathway, became significant. Among the pathways that lost significance between the two LMMs there were pathways like neurotransmitter receptors and postsynaptic signal transmission (Reactome), Parkinson’s Disease (KEGG), Oxidative Phosphorylation (KEGG), and alpha-synuclein (PID) pathways.

**Figure 3:**
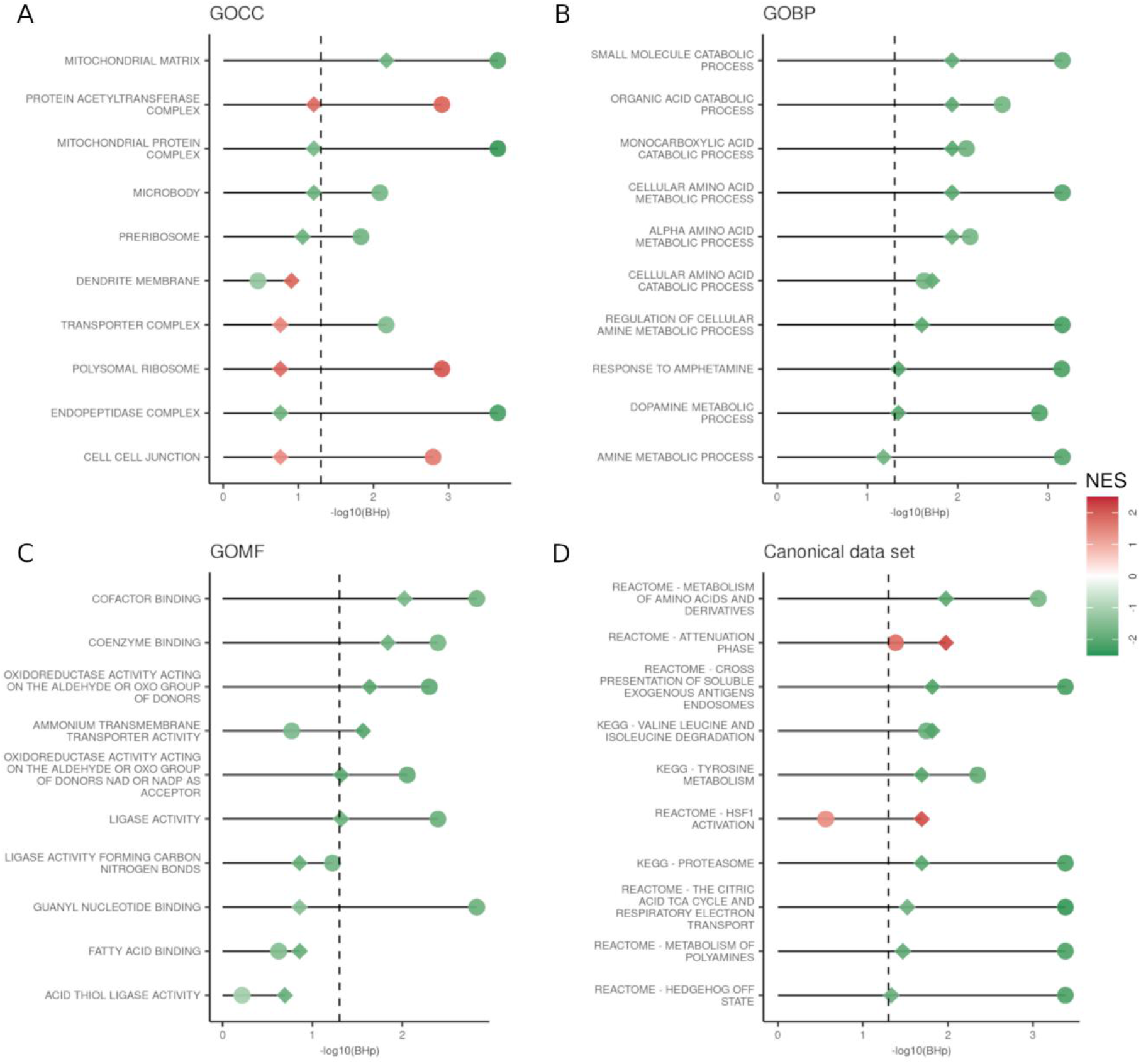
Comparison of the GSEA for the gene ranked by two linear mixed models (LMMs). Top 10 significant hits of the cell-proportions-aware (second) LMM (diamond) compared with their respective values from the cell-proportions-unaware (first) LMM (circle). Each shape is colored by the normalized enriched scores (NES), -log10 of the adjusted p-value is reported on the x-axis. Significance threshold (BHp=0.05) is indicated with by the dashed vertical line.

**Figure 4:**
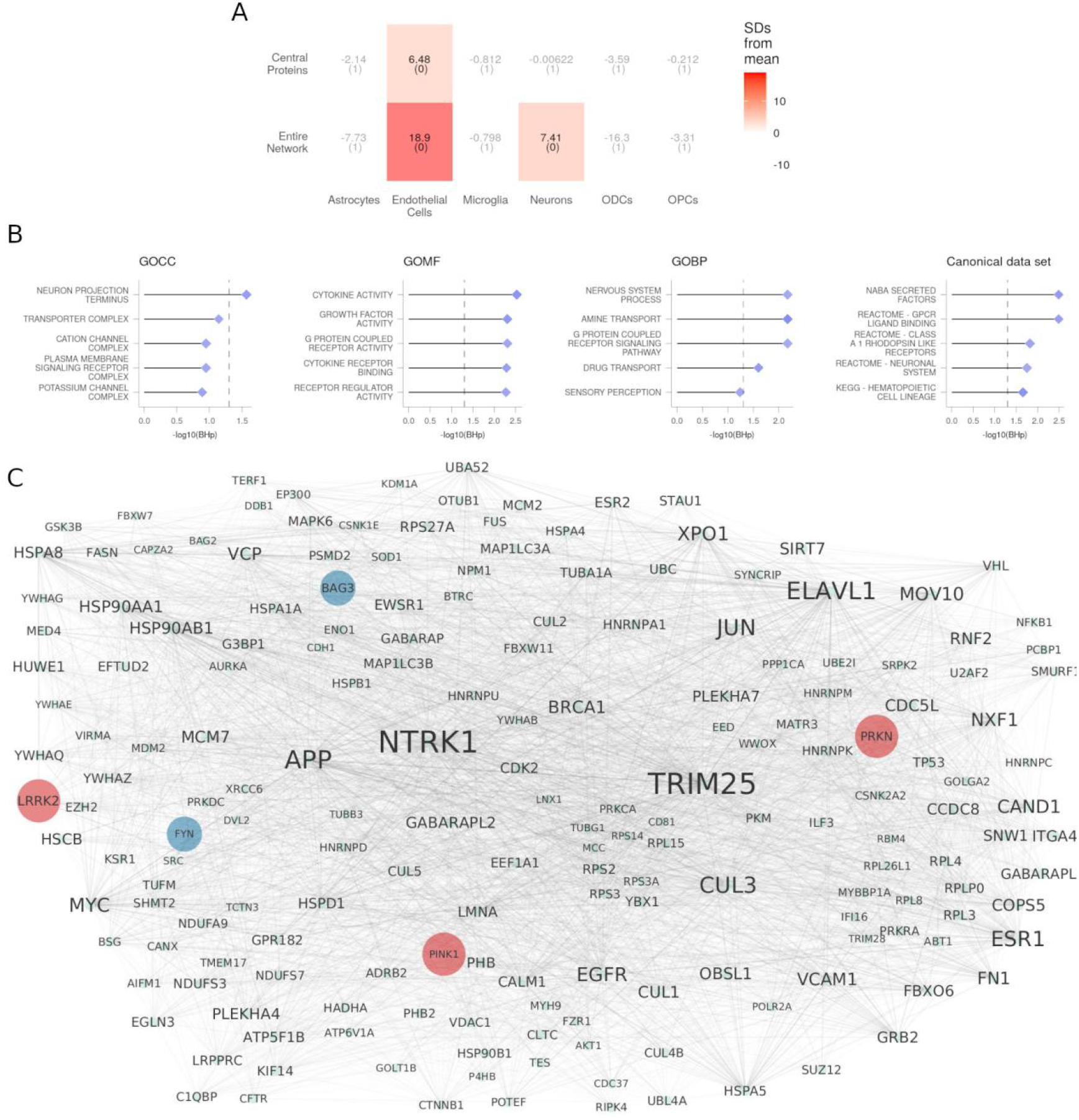
PPI network analyses. **A**. EWCE results for the full network and central protein. The number of standard deviations of the expression of each list from the mean level expression of 10.000 equally sized random lists is reported for each cell type. BH-corrected p-values are reported between brackets. **B**. GSEA results of the nodes in the network ranked by betweenness centrality. -log10 of the adjusted p-value is reported on the x-axis and significance threshold (BHp=0.05) is indicated by the dashed vertical line. **C**. Network of the central proteins extracted from the full PPI network. Label size is proportional to the vertex degree, edge thickness and color proportional to the edge betweenness. PD-causing genes (LRRK2, PINK1, PRKN) and the genes closest to a risk factor (BAG3, FYN) are highlighted respectively with brick red and cerulean blue.

### 4. PPI partners of DEGs are enriched for genes expressed in endothelial cells and neurons

To identify the interaction partners of the proteins encoded by the DEGs, we constructed a protein-protein interaction (PPI) network based on the BioPlex, Biogrid, HuRI, IID, IntAct, and String databases (Methods). Starting with the 100 DEGs, we obtained a network comprised of 5,615 vertices and 92,035 edges (Methods, Extended Data Table 3-1). We assessed the expression-weighted cell-type enrichment with EWCE of the entire network and found a significant enrichment for endothelial cells and neurons (Figure 3A, Extended Data Table 3-2). Enriched pathways (using GSEA) were secreted factors, G-protein-coupled receptors, amine transport, catecholamine secretion, neuronal system, and hematopoietic cell lineage differentiation (Figure 3B, Extended Data Table 3-3). Moreover, this network was significantly enriched in PD-causing genes (6 genes in the network out of 6, FET p-value 4.6*10^−4^) and in genes in proximity of the GWAS risk variants (36 genes in the network out of 97, FET p-value 0.029).

### 5. Central proteins of DEG PPI partners are known PD-causing genes and genes in proximity to PD risk factors

Within the PPI network of DEG interaction partners, we identified 198 nodes that have both a degree centrality higher than the 95^th^ percentile (276 nodes) as well as a betweenness centrality higher than the 95^th^ percentile (281 nodes). Among these, 11 (6%) were encoded by DEGs, hence biased towards higher centrality. The remaining 187 central nodes were defined as top central proteins (Figure 3C, Extended Data Table 3-1). EWCE analysis revealed that these genes are enriched for endothelial cells (Figure 3A, Extended Data Table 3-2). The top proteins were also enriched in PD-causing genes (3 genes out of 6, FET p-value 6.8*10^−4^) and included Leucine-Rich Repeat Kinase 2 (*LRRK2*), Parkin RBR E3 Ubiquitin Protein Ligase (*PRKN*), and PTEN-induced putative kinase 1 (*PINK1*). Moreover, several of the genes reported in proximity to GWAS risk factors were among the central proteins too: LRRK2, BAG Cochaperone 3 (*BAG3*), and FYN Proto-Oncogene and Src Family Tyrosine Kinase (*FYN*).

## Discussion

We analyzed the influence of cyto-architectural alterations on the transcriptomic signals from human *substantia nigra* microarray data from PD patients and CTRL. We demonstrated that a broad palette of alterations in cell composition was present within and between strata as well as across studies. Specifically, a significant decrease of neurons and increase of ODCs in PD with respect to CTRL were the most consistent, but also differences for microglia and OPCs were found. The lack of a universal alteration pattern might be attributed to the mixed cohort of patients assessed, with variable putative etiological factors, phenotype, and disease severity that characterize PD, as well as technical variability. Nonetheless, a meta-analysis of cell proportions showed a significant decrease in the neurons and an increase in ODCs and endothelial cells in PD. This pervasive heterogeneity heavily influenced differential expression analysis between PD patients and controls. When not correcting for cell type composition we found 1,072 DEGs in a meta-analysis. After adjusting for cell type composition, only 67 DEGs remained, next to 33 new DEGs. Together, these findings provided evidence that the systematic integration of microarrays of *substantia nigra* in PD, albeit a popular methodology to increase power in detecting the DEGs, resulted in many spurious associations when not controlling for cyto-architecture. The DEGs identified with the second model showed a down-regulated signature in neurons and an up-regulated one in the OPCs adding to the mounting evidence for the involvement of neurons and oligodendrocyte lineage cells in PD (Reynolds et al., 2019; Agarwal et al., 2020; Bryois et al., 2020).

Estimated cell-type proportions were supported by previous and independent studies confirming the reliability of the deconvolution strategy. Firstly, the observed loss of dopaminergic neurons in *substantia nigra* is a pathological hallmark of PD. Secondly, our observation that endothelial cells were increased in the *substatia nigra* of PD patients is in line with previous reports (Faucheux et al., 1999; Desai Bradaric et al., 2012), although, it is still unclear whether endothelial cell expansion is a result or a driver of the inflammation status. Indeed, angiogenesis can be stimulated by molecules secreted by astrocytes and microglia in the reactive status (Naldini and Carraro, 2005; Wada et al., 2006) in a vicious loop (Barcia et al., 2004). Thirdly, the increase in microglia in one of the studies can be related to reactive gliosis present in PD that is known to implicate this cell type (Vila et al., 2001). Finally, the observed increase in OPCs and ODCs reflects the skewed neuron/oligodendrocyte ratio in the dissected samples due to the neuronal death.

The deconvoluted cellular proportions did not only uncover interesting features of the cyto-architecture of the *substantia nigra* in PD and adjusted the DEGs detection but also influenced the pathway analysis. We confirmed alterations in known pathways and functions like the tricarboxilyc acid cycle (Grünewald et al., 2019), the dopamine metabolism (Masato et al., 2019), the proteasome activity (McNaught et al., 2003), the HSF1 activation and attenuation (Gomez-Pastor, 2018), and the expression of genes activated in the hedgehog pathway in the off status (Gonzalez-Reyes et al., 2012). Furthermore, we also identified an intriguing decrease in the ammonium transport proteins which, despite having been suggested for other pathologies (Adlimoghaddam et al., 2017), has not yet been reported for PD. Intriguingly, previous data suggest that ammonia accumulation could affect energy metabolism, mitochondria, inflammation, and neurotransmission (Cooper and Plum, 1987; Kelly and Rose, 2010). Importantly, accounting for the cyto-architecture also showed that several of the usually reported pathways (e.g. Parkinson’s disease pathway, Oxidative Phosphorylation, alpha-synuclein pathway) might be driven, at least in part, by changes in cell composition rather than the pathological status. A similar observation has recently been reported also in PD prefrontal cortex (Nido et al, 2020), reinforcing the contention that cyto-architecture is an important covariate with a major impact on our ability to understand transcriptional changes in bulk transcriptomics.

As interacting partners of DEGs can have consequent altered functionality, we constructed a network of protein-protein interactors with the detected DEGs. Cell type enrichment analysis of this network corroborated that gene deregulation might have an impact on neuronal biology also through the interacting partners of the DEGs. Further, it showed ramifications in endothelial cell processes unidentified by the differential expression analysis. Similarly, the GSEA reinforced the involvement of the neurons and pinpointed to previously undetected immune response terms. Importantly, PD-causing and genes proximal to GWAS variants, albeit not being differentially expressed, were enriched in this network. This convergent evidence shows how genetically-identified genes can have an impact on the transcriptional landscape of substantia nigra even when not differentially expressed and supports the relevance of this network. Exploiting their topological characteristics, we nominated key proteins that were central in this partner network. Some of the central proteins have been indeed implicated in the pathogenesis of PD by independent lines of research, such as genes whose variants are causative of inherited forms of PD (*LRRK2, PINK1, PRKN*) (Kitada et al., 1998, Tolosa et al., 2020, Valente et al., 2004), and/or are the closest to risk variants for developing PD (*BAG3, FYN, LRRK2*) (Nalls et al., 2019). Furthermore, other hits with compelling evidence supporting their role in neurodegeneration (e.g. *GSK3β, WWOX*, and VPC; Chan et al., 2012, Wang et al., 2012) corroborate that our approach can be used to identify new potential players in the PD pathogenesis (e.g. *NTRK1, TRIM25, ELAVL1*).

Our cell-type composition aware meta-analysis comes with some limitations. Firstly, the deconvolution step does not take into account the differences in cell size or in RNA content of the various cell types potentially leading to systematic errors (Zaitsev et al., 2019). Secondly, as the sum of the estimates of the cell-type proportions is constrained to be 100% for each sample, an increase or decrease in any of the cell types thus has to result in an equal but opposite alteration in at least one of the other cell types. Thirdly, the incomplete annotation of the samples across the studies prevented us from exploring the effect of the age, age of onset, pathology progression, genotype, and Braak staging on the cell proportions and transcriptional processes, all known to influence PD severity (Pagano et al., 2016; Fereshtehnejad et al., 2017; Koros et al., 2017). The future advent of scRNAseq/snRNAseq studies of human *substantia nigra* of PD patients will allow additional steps forward in this area of research.

In conclusion, our meta-analysis of bulk transcriptomics gives an updated view of the transcriptional landscape in the *substantia nigra* of PD patients. In addition to leverage a big number of studies and samples similarly to previous works (Zheng et al., 2010; Wang et al., 2019a), we accounted also for the dramatic cyto-architectural alteration induced by PD in this brain area uncovering its effects on downstream analyses. Using multiple complimentary approaches, encompassing the transcriptional and protein-interaction perspectives, we implicate neurons, OPCs, and endothelial cells as affected in PD *substantia nigra*. Moreover, a number of novel PD candidates and the identified enriched biological processes offer clues for a better understanding of the complex PD pathology, and provide steppingstones for the identification of new therapeutic approaches.

## Supporting information

EDT2-1

EDT2-2

EDT2-3

EDT2-4

EDT3-1

EDT4-1

EDT4-2

EDT4-3

EDF2-1.tif

EDF2-2.tif

## Extended Data Legends

**Extended Data Figure 2-1:** *Differential expression analyses extended figures*. ***A***. *Spearman’s correlation between the cell estimates across the studies*. ***B***. *Violin plots showing the level of expression of representative genes in the microarrays before and after correcting for gender only (orginal model) in red, or for gender and the cell estimates (corrected model) in blue*. CHD9 *(cell-unaware-LMM β 0*.*229±0*.*047 SD BHp 0*.*0002; cell-aware-LMM β 0,156±0*.*052 SD BHp 0*.*109), and* CLK4 *(cell-unaware-LMM β 0*.*184±0*.*043 SD BHp 0*.*001; cell-aware-LMM β 0*.*107±0*.*043 SD BHp 0*.*199), two DEGs of the cell-unaware-LMM only;* OXTR *(cell-unaware-LMM β −0*.*174±0*.*075 SD BHp 0*.*097; cell-aware-LMM β −0*.*32±0*.*076 SD BHp 0*.*013) and* SYN2 *(cell-unaware-LMM β −0*.*023±0*.*056 SD BHp 0*.*844; cell-aware-LMM β 0*.*215±0*.*046 SD BHp 0*.*004), two DEGs of the cell-aware-LMM only*. ***C***. *Heatmap showing the expression of the DEGs from the cell-proportions-aware LMM in the cell types identified in GSE140231. Gene expression values have been scaled for visualization purposes*.

**Extended Data Figure 2-2:** *Pearson’s correlation matrices of the expression of the top 20 selected markers per microarray data set from the BRETIGEA package and the human substantia nigra snRNAseq (GSE140231)*.

**Extended Data Table 2-1:** Results of the linear model to test the differences in the estimates of the cell types between CTRL and PD.

**Extended Data Table 2-2:** Results of the metafor meta-analysis of the cell estimates across the studies.

**Extended Data Table 2-3:** *DEGs identified with the two LMMs. Sheet 1: DEGs identified with the cell-proportions-unaware model. Sheet 2: DEGs identified with the cell-proportions-aware model*.

**Extended Data Table 2-4:** *Results of the EWCE of the up-regulated and down-regulated genes from the cell-proportions-aware LMM in the GSE140231 summary expression matrix*.

**Extended Data Table 3-1:** *GSEA of the genes ordered by the signed p-values from the two LMMs. Sheet 1: GOCC. Sheet 2: GOMF. Sheet 3: GOBP. Sheet 4: Canonical pathways collection*.

**Extended Data Table 4-1:** *Network metrics. Nodes betweenness centrality, degree centrality, and ranking by the two metrics. Hubs, bottlenecks, and central protein lists*.

**Extended Data Table 4-2:** *Results of EWCE for the nodes in the network and the central proteins*.

**Extended Data Table 4-3:** PPI *GSEA results. Sheet 1: GOCC. Sheet 2: GOMF. Sheet 3: GOBP. Sheet 4: Canonical pathways collection*.

