## Supplementary figures and images for "Correcting differential gene expression analysis for cyto-architectural alterations in substantia nigra of Parkinson’s disease patients reveals known and potential novel disease-associated genes and pathways"

### EDF2-1.tif

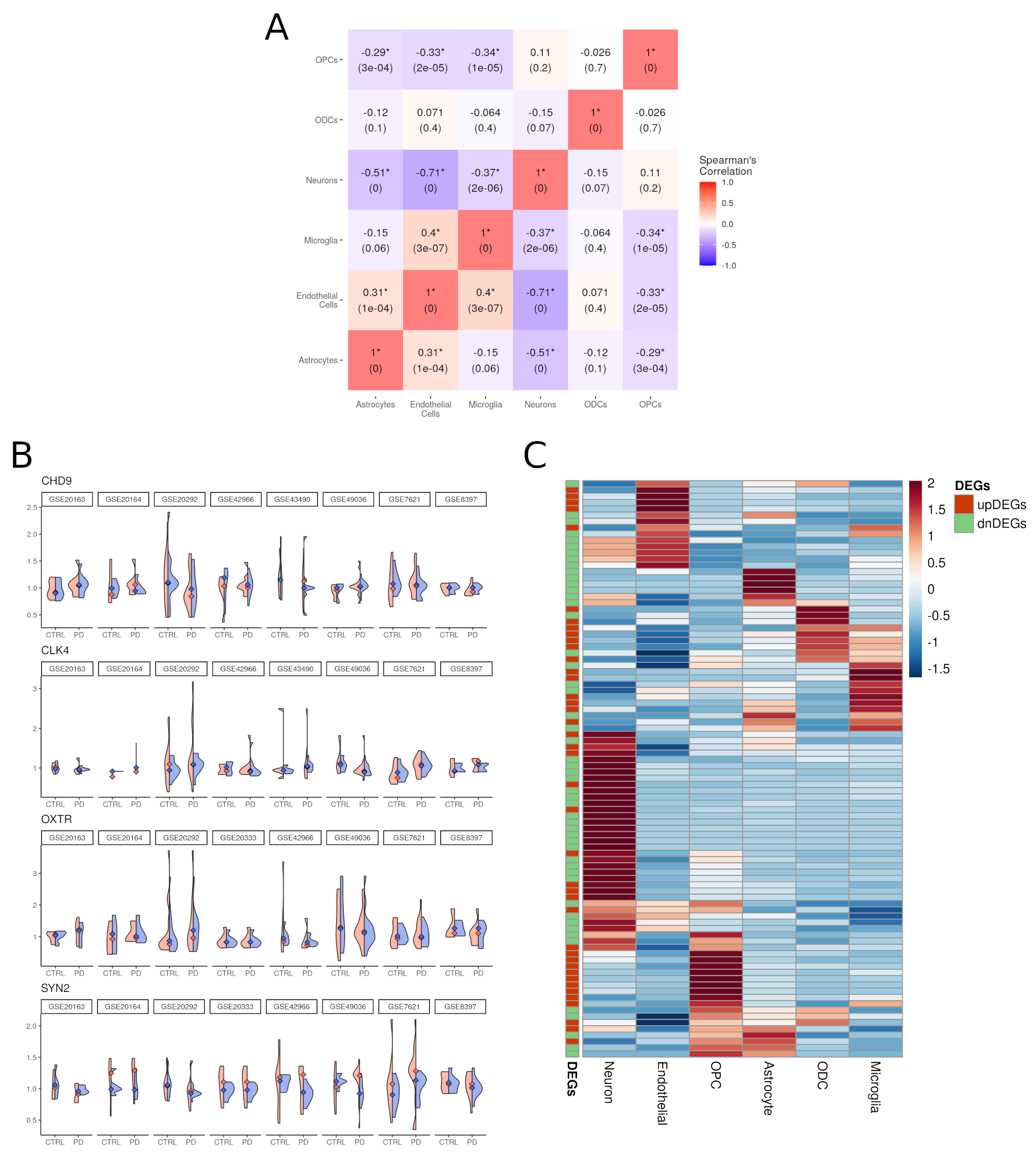

### EDF2-2.tif

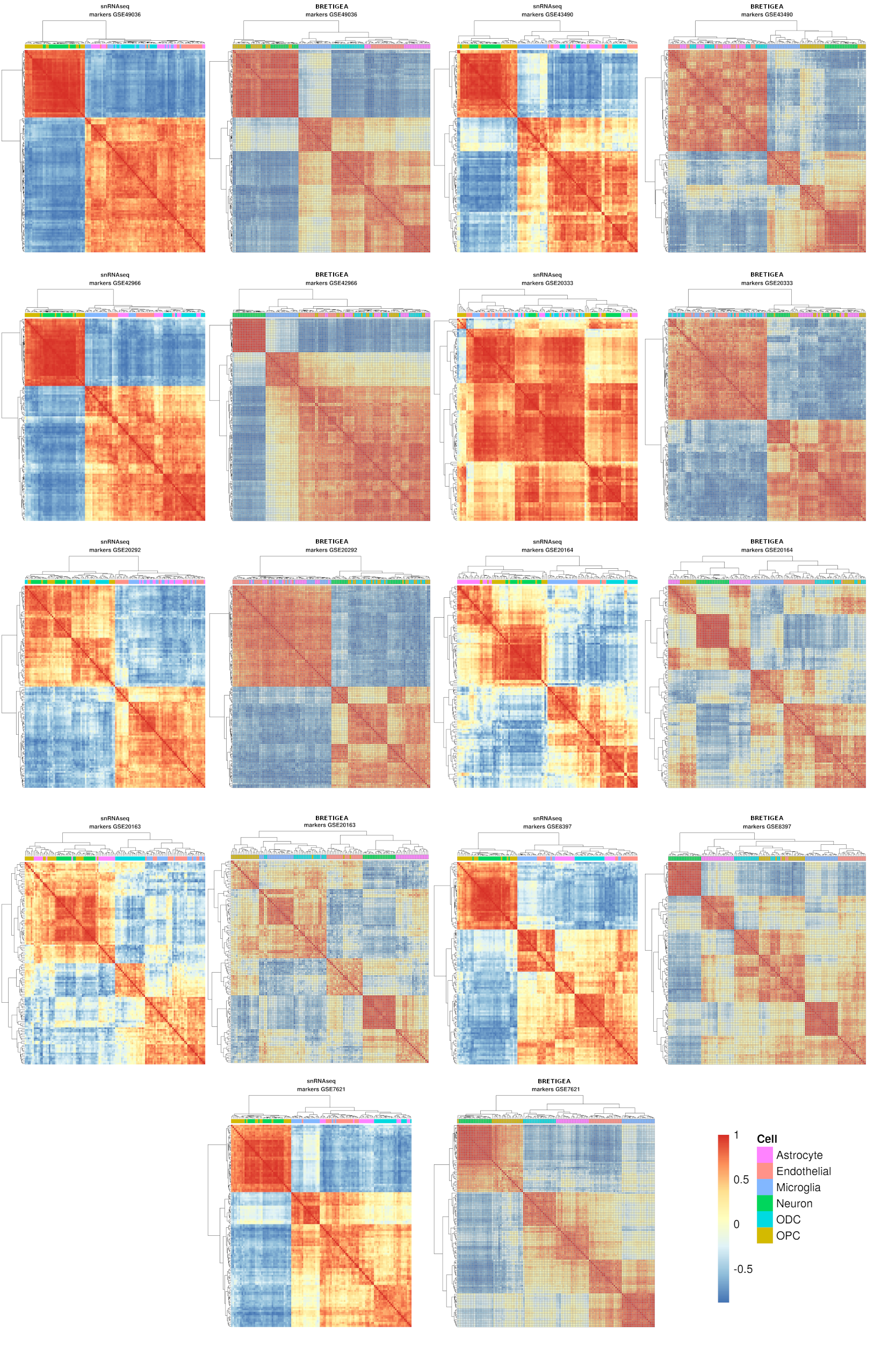
